# Homology modeling in a dynamical world

**DOI:** 10.1101/135004

**Authors:** Alexander Miguel Monzon, Diego Javier Zea, Cristina Marino-Buslje, Gustavo Parisi

## Abstract

A key concept in template-based modeling is the high correlation between sequence and structural divergence, with the practical consequence that homologous proteins that are similar at the sequence level will also be similar at the structural level. However, conformational diversity of the native state will reduce the correlation between structural and sequence divergence, because structural variation can appear without sequence diversity.

In this work, we explore the impact that conformational diversity has on the relationship between structural and sequence divergence. We find that the extent of conformational diversity can be as high as the maximum structural divergence among families. Also, as expected, conformational diversity impairs the well-established correlation between sequence and structural divergence, which is nosier than previously suggested. However, we found that this noise can be resolved using *a priori* information coming from the structure-function relationship. We show that protein families with low conformational diversity show a well-correlated relationship between sequence and structural divergence, which is severely reduced in proteins with larger conformational diversity. This lack of correlation could impair Template-based modelling (TMB) results in highly dynamical proteins. Finally, we also find that the presence of order/disorder can provide useful beforehand information for better TBM performance.

**Author summary:** Template-based modelling (TBM) is the most reliable and fastest approach to obtain protein structural models. TBM relies in the high correlation between sequence and structural divergence, with the practical consequence that proteins that are similar at the sequence level will also be similar at the structural level, allowing in this way the selection of the better template to obtain the 3D model of the target sequence. However, protein native state could be described by a collection of conformers in equilibrium where their structural differences are called conformational diversity.

In this work, we explore the impact that conformational diversity has on the relationship between structural and sequence divergence. We firstly found that the extent of conformational diversity can be as high as the maximum structural differences reached by families differing in their sequences. In these proteins with higher conformational diversity levels, the well-established correlation between sequence and structural divergence is nosier than previously suggested due to the presence of structural change without sequence variation. This lack of correlation could impair TBM results due to the uncertainty in the correct template selection. Finally, we also found that the presence of order/disorder can provide useful beforehand information for better TBM performance.

## Introduction

Template-based modeling (TBM) is the most reliable, accurate, and fastest approach for protein structure prediction[1–3]. The accumulation of experimental structures in the Protein Data Bank (PDB) has increased the fold-space coverage[4], which in combination with the steady enhancement of template-detection techniques over the last several years[2], allows prediction of 3D structures in at least 50% of the human proteome, and almost 70% for some prokaryotic proteomes using current TBM methods[5, 6]. TBM relies on the fact that homologous proteins, with detectable sequence similarity, possess similar 3D structures. Pioneering work by Chothia and Lesk found that structural divergence (SD) increases with evolutionary distance, measured as percent identity, following a non-linear relationship[7]. Very similar sequences show modest structural differences, which suddenly increase when sequence identity drops below 30%. This trend was later confirmed by others[8–10]; however, in recent studies a linear correlation between sequence and structural divergence has been found[11–13].

Using the relationship between sequence identity and structural distance, the first step in TBM involves the search for an adequate template[3]. In view of the above-mentioned studies on structure-sequence relationships, the better template will certainly be one with the maximum sequence similarity to the target sequence in the known structural database. Evolutionary distances between target and template, the presence of ligands, and resolution are also useful guidelines for template selection[14]. This step is followed by alignment between the template and target sequence to detect conserved and variable regions. The final step of TBM is refinement through a combination of methods to render a 3D model of the target sequence. Using sequence identity as a measure of the distance between target and template sequences, it was found that structural models differ 1-2 Å C-alpha root mean squared deviation (RMSD) from a selected native structure for templates with more than 50% sequence identity. In the case of templates between 30-50% identity, the distance between a model and a native structure is about 4 Å RMSD, while for templates below 30% identity, template-free methods outperform TBM techniques[1].

In spite of the outstanding contributions of TBM approaches to a great variety of fields[15], it is still difficult to obtain high quality 3D models. Errors derived from target and template alignments[16], along with refinement of the initial model to obtain more native-like models[14], are among the major problems to solve in order to improve 3D model quality[3]. However, there is still a conceptual issue to face in order to improve TBM predictions. This issue is related to the nature of the “native state” of proteins, which are composed of different conformers in equilibrium, a key concept for understanding protein function[17]. In this sense, TBM techniques should progress towards a new step in its development to predict the “native state of proteins”, and not simply to “predict the structure” (in terms of the alpha carbon scaffold) of a target sequence. Several authors have previously pointed out the impact of conformational diversity on TBM approaches[18], primarily because a given template (with a determined distance to the target sequence) can have different conformers sampling a large conformational space[12, 19]. A wide range of structural differences among conformers can be observed by comparing structures of the same protein obtained under different crystallization conditions. These differences result from the relative movements of large domains [20], secondary and tertiary element rearrangements[21], and loop movements[22], which overall can produce a conformational diversity up to 4-5 Å of RMSD [23–25]. Even up to 15-20 Å can be observed, depending on the structural alignment algorithm used to calculate the RMSD[18]. Taking into account this extent of conformational diversity, performance of TBM methods should be re-evaluated. Blind evaluation protocols use only one conformation of the selected templates, and the performance of the resulting model depends highly on that selection[26].

Underneath the effect of conformational diversity in TBM techniques, the more complex problem of solving how structural information is codified in the protein primary structure remains[13]. The so-called local model maintains that a few positions in the protein define the global structural arrangement. Non-linear behaviour in the structure-sequence relationship supports this hypothesis due to the observation that a large amount of sequence variation is required in order to dramatically change the structure (mostly below 20-25% sequence identity). On the contrary, the global model supports the idea that several positions spread along the protein define the structural arrangement. A linear relationship between structural change and sequence divergence will support this model by showing proportional change between those variables. However, considering that a single sequence can adopt several conformations, makes it even more complicated to predict how non-synonymous substitutions correlate with structural divergence.

As a key concept to be taken into account in TBM approaches, here we explore the impact of conformational diversity on the relationship between structural and sequence divergence. To this end, a curated data set of 2024 proteins with experimentally known conformational diversity, clustered in 524 homologous families (>30% local identity and 90% coverage), were analysed to derive structural and sequence similarities. We found that the use of a highly redundant sequence dataset (that is, considering the conformational diversity) blurs the well-established relationship between sequence and structure divergence more than shown in previous studies. However, we also found that this trend could be solved using *a priori* information from the structure-function relationship. We show that families containing proteins with low conformational diversity, which we call “rigid” proteins, show a well-correlated behaviour of sequence and structural divergence; on the contrary, this correlation is severely reduced in protein families with larger conformational diversity. This lack of correlation could impair TBM results in highly dynamic proteins. Finally, we found that the presence of order/disorder regions could be useful prior information resulting in a better TBM performance.

## Results

### Protein conformational diversity can be as high as the structural divergence in family evolution

We performed an “all against all” structural alignment within each family using MAMMOTH[27], against our dataset of 524 protein families containing 2024 proteins with known conformational diversity, totalling 37755 structures (at least 5 conformers of each protein, with ~19 conformers in average) extracted from the CoDNaS database[23]. For each pair of homologous proteins within each family, the percent sequence identity was calculated, aligning each pair of sequences with the Needleman-Wunch algorithm[28]. Since each protein is represented by an ensemble of conformers, the maximum RMSD derived from an “all vs all” comparison of conformers, belonging to the homologous proteins being compared, is called maximum structural divergence (hereafter MSD for simplicity). When this procedure is repeated between the pairs of conformers (different structures of the same protein, see Methods), the maximum conformational diversity (hereafter CD for simplicity) is obtained, measured as the maximum RMSD of a given protein (see schematic protocol in **S1 Fig**). In **S2 Fig,** the plot contains all comparisons between conformers for each pair of homologous protein being studied, containing ~3.5 million dots. **Fig 1a** shows the relationship between the MSD (green points) versus percent sequence identity. **Fig 1a** also shows the CD values (red dots). Green dots show the typical behaviour previously found between sequence and structure divergence[7, 13] with a steep decrease of structural similarity at low identity percent (below ~30%). However, our dataset also shows high SD at very high sequence similarities, as a consequence of proteins with conformational diversity as high as the structural divergence of the family.

**Fig 1.**
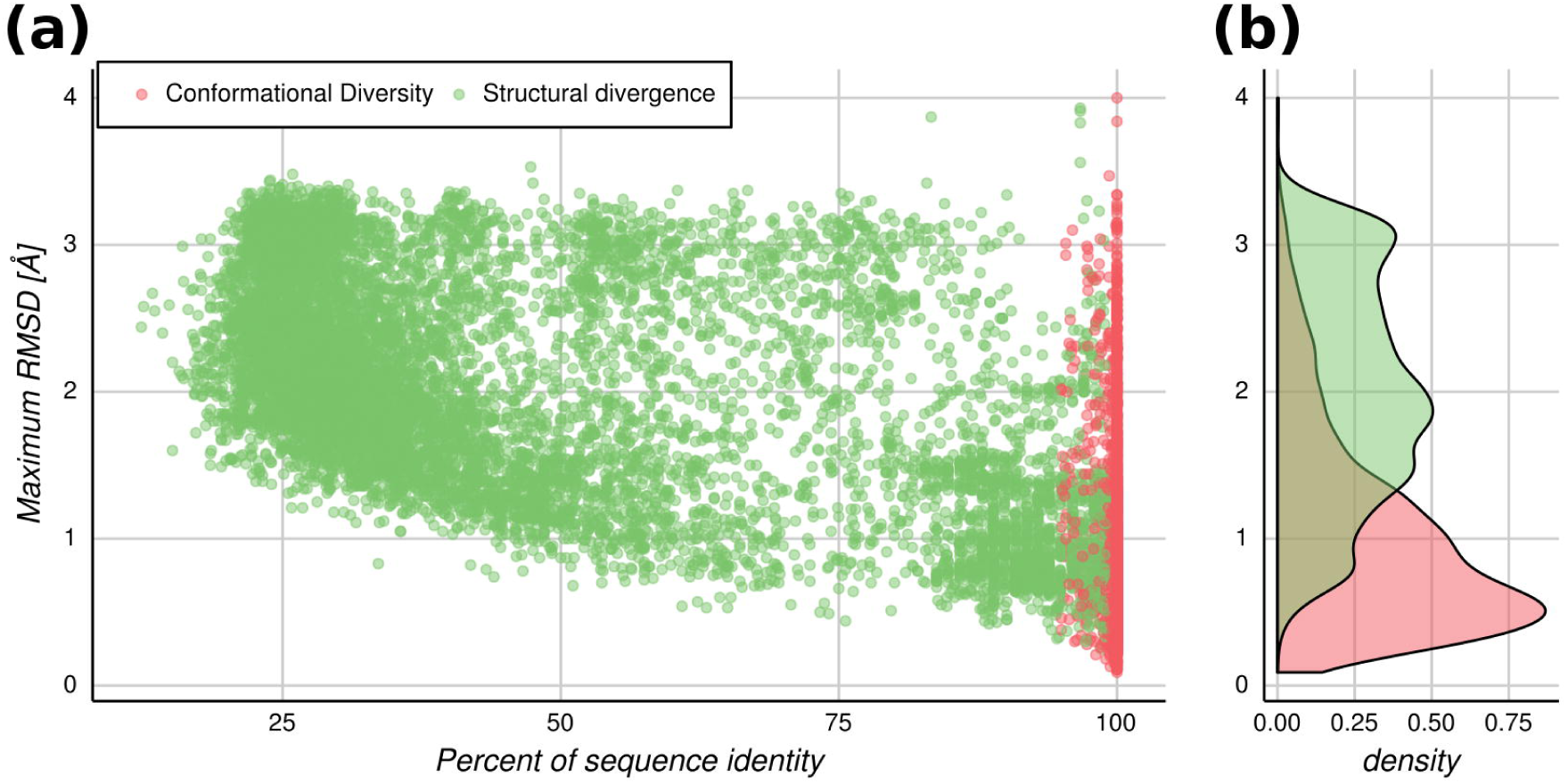
Maximum RMSD (MSD and CD) versus sequence percent identity. Points refer to the maximum RMSD obtained from an “all vs all” comparison between structures from two homologous proteins (MSD), or from the same protein (CD). (a) Green dots: comparisons between homologous protein pairs. Red dots: comparison between conformers of the same protein. (b) Distributions of the maximum RMSD between two homologous proteins (green) and between conformers of the same protein (red).

The distribution of the CD shows (**Fig 1b**) mostly moderated RMSD values, with an average of 1 Å. Nevertheless, it also shows high positive skewness, towards larger RMSD values. This is in concordance with previous work[18]. The 90th percentile of the analysed proteins show a CD below 2 Å of RMSD, and then 10% of the proteins can exist in a conformational space as large as the MSD, coming from comparison of remote homologous proteins (~3 Å). This is an interesting result that indicates that a given sequence can potentially exist in a conformational space as big as the structural divergence that arose from the accumulation of substitutions, namely the evolutionary process. In the light of the conformational diversity, it is easy to understand that closely homologous proteins (suppose above 80% sequence identity) can have either high or low RMSD values when superposing their structures, depending on the particular conformers being compared. So, conformational diversity can lead to large RMSD values between proteins over short evolutionary time periods, instead of reaching these RMSD values through the long process of accumulation of sequence mutations.

The other important consequence of this finding is that the correlation between sequence and structural divergence is weaker than stated in previous works[11–13]. In other words, due to the CD, a given sequence can adopt different conformations, so the structural change due to non-synonymous substitutions in a divergent evolution process will make the relationship between sequence and structure noisier. In fact, the distribution of the MSD (green dots in **Fig 1a**) and sequence identity, have a Spearman’s rank correlation rho of -0.52. The relationship between sequence and structure will be visible in light of the conformational diversity, as explained below.

### Template selection and structural diversity

As mentioned before, TMB approaches require the use of a protein with known structure as the template. This identification can be performed using a broad variety of techniques with different sensitivities[3,29,30]. A key point in this step is selection of the best template, which is based upon the commonly used relationship between structural and sequence divergence[7], and the one maximizing both coverage and percent sequence identity against the target[31]. However, as we can see from **Fig 1,** this criterion is not as simple as previously stablished. In **Fig 2,** we show the MSD distribution among bins of 10% sequence identity between pairs of homologous proteins. It is possible to observe the great variation in RMSD for each particular bin. More importantly, the maximum RMSD value is almost equal for all the considered bins of percent identity (mean = 3.54 Å and standard deviation = 0.23 Å). Therefore, template selection is not as straightforward as just selecting a structure in a given identity bin, because it is not known how those structures belong to the conformational ensemble of the sequence to be modeled[26].

**Fig 2.**
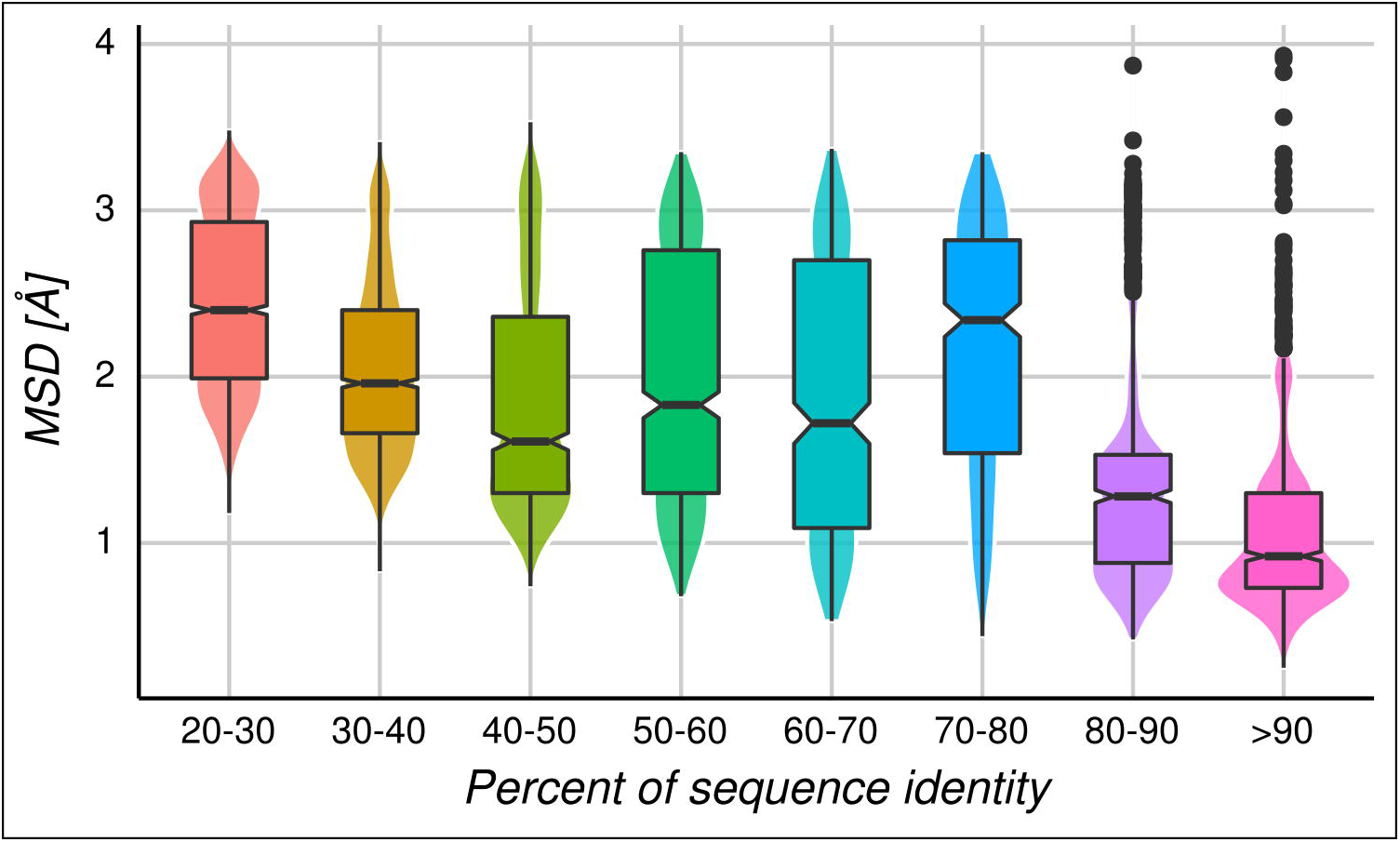
MSD distributions over all homologous protein pairs by bins of 10% sequence identity. Above and below the median (horizontal line inside each box) are the first and third quartiles, respectively. The notches display the median absolute deviation (M.A.D).

However, the distributions per bin shown in **Fig 2** could be influenced by the presence of a given protein family with exceptionally large or small structural diversity. For that reason, in **Fig 3** we show the average MSD per protein family in a bin of sequence identity. The averaged RMSD values are between 1.34 and 2.10 Å for the different bins, with standard deviations between 0.19 and 0.29, showing that the dispersion is not related to the sequence identity. In **S3 Fig,** we show that the average MSD does not depend on how populated the corresponding family is.

**Fig 3.**
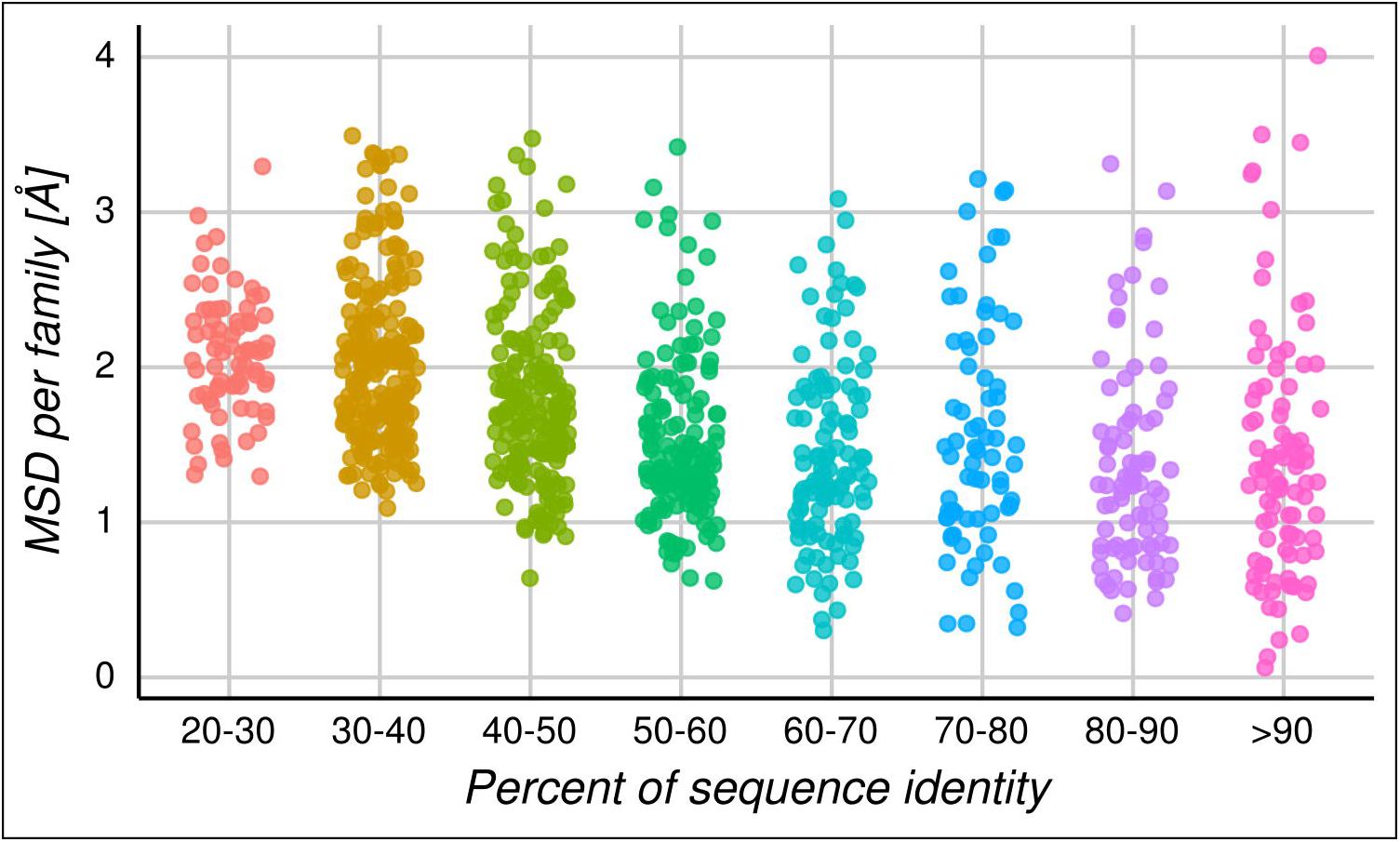
MSD distributions over bins of 10% sequence identity per family. Each dot represents the average MSD for all homologous protein pairs per family in a given bin of percent identity. It is possible to see a great dispersion of structural divergence even at low percent identity and that the different families spread about ~2.9 Å RMSD in average per bin.

Taking these results into account, the selection of an adequate template will depend heavily on the target protein, whether it has a high or a small conformational diversity. Which then will be the general recommendations for selecting a good template? The relationship between SD and CD will give us a clue.

### How does structural divergence correlate with conformational diversity?

In a simple evolutionary scenario of the evolution of a protein family, we can consider that CD is a conserved trait among its members and that the common ancestor of the family shows low conformational diversity at the backbone level. If all the conformers belonging to this common ancestor are structurally aligned, the resulting RMSD values would be about 0.5 Å (a value equivalent to the crystallographic error), meaning that the conformer population is almost identical. It is important to say that these conformers are structurally equivalent at the backbone level, as the RMSD is measured using the alpha carbons, but conformational differences at the residue level cannot be discarded[32, 33]. Considering that this family has selective pressure to maintain its conformational diversity in most of their proteins (i.e. due to functional restrictions[34]), most of the structural divergence of this family will have originated by the accumulation of nonsynonymous substitutions. On the contrary, in the case that a family originated with a common ancestor showing large conformational diversity (i.e. a RMSD of ~2 Å), the process of divergence due to the accumulation of nonsynonymous substitutions will certainly increase the available repertory of conformations and eventually increase the structural divergence. We have found that the dispersion of the CD extent is rather low, possibly indicating that the CD can be a trait conserved within families (See **S4 Fig**).

It is our central hypothesis that the maximum conformational diversity of a protein will correlate with the maximum SD that can be reached by that family. To probe this hypothesis, the average of the CD and the MSD per family was calculated, and a Pearson’s correlation coefficient of 0.75 (P-value < 0.01) was observed (see **Fig 4**).

**Fig 4.**
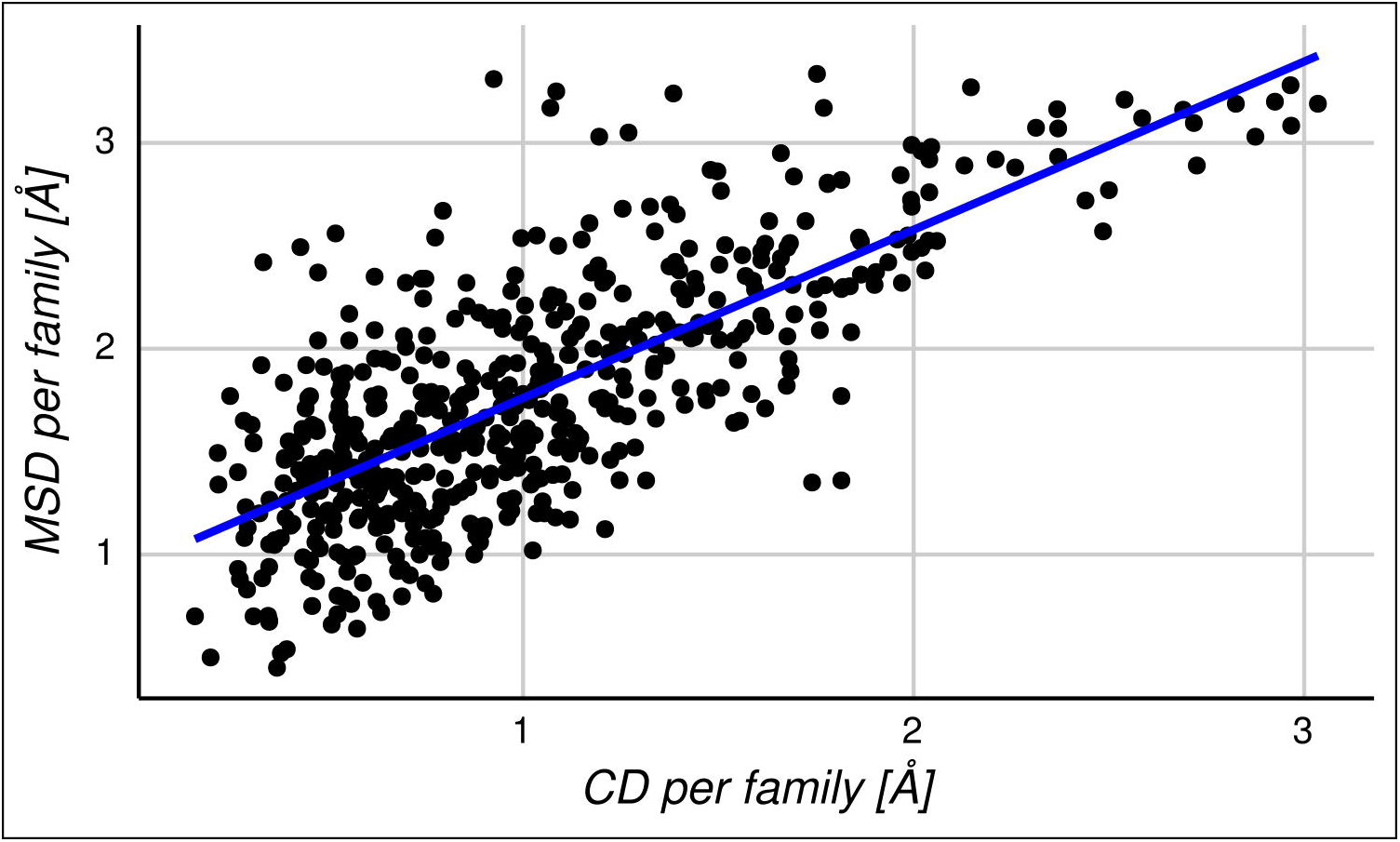
Relationship between MSD and CD. Each dot represents the average RMSD values for the MSD and the CD in a specific family. The data show a Pearson’s correlation coefficient of 0.75.

As MSD could change proportionally with the sequence divergence of each family, in **S5 Fig** we show that the association is independent of the sequence divergence found in each family. In the light of the results shown in **Fig 4**, we study the relationship between structural and sequence divergence by splitting the dataset into homologous protein pairs with large and small conformational diversity (>= 0.5 Å and < 0.5 Å RMSD respectively, obtained as the average of CD between each protein). This threshold is near the crystallographic error (~0.5 Å [35]). It is interesting to note that the Spearman’s rank correlation rho between structure (MSD) and sequence divergence (sequence identity percent) is -0.83 (with a significant P-value < 0.01) in the subset of protein families with small conformational diversity per homologous protein pair, and -0.51 (P-value < 0.01) in the subset of protein families with large conformational diversity (see **Fig 5**). These results indicate that the known correlation between sequence and structure[7] is strong in the subset of protein families with low conformational diversity. In these families, the biological function can be achieved with conformers almost identical at the backbone level. That will make the relationship between sequence and structure straightforward, namely, the change in structure is proportional to the observed change at the sequence level. In the opposite cases, for the subset of families with high conformational diversity, two scenarios are possible: either the biological function is less tight with a single conformation, or inversely, the function requires high plasticity of the structure. In this sense, the subset with conformational diversity below 0.5 Å RMSD has a linear and exponential fit with R^2^ values of 0.54 ± 0.15 and 0.66 ± 0.12, respectively. While for the subset with conformational diversity above 0.5 Å, RMSD has a linear and exponential fit with R^2^ values of 0.23 ± 0.21 and 0.29 ± 0.18, respectively. These R^2^ values are the mean obtained for testing datasets in a 5-fold cross validation. We also found that splitting the distribution in bins of CD, these correlation coefficients change monotonically as the CD increases (**S1 Table**).

**Fig 5.**
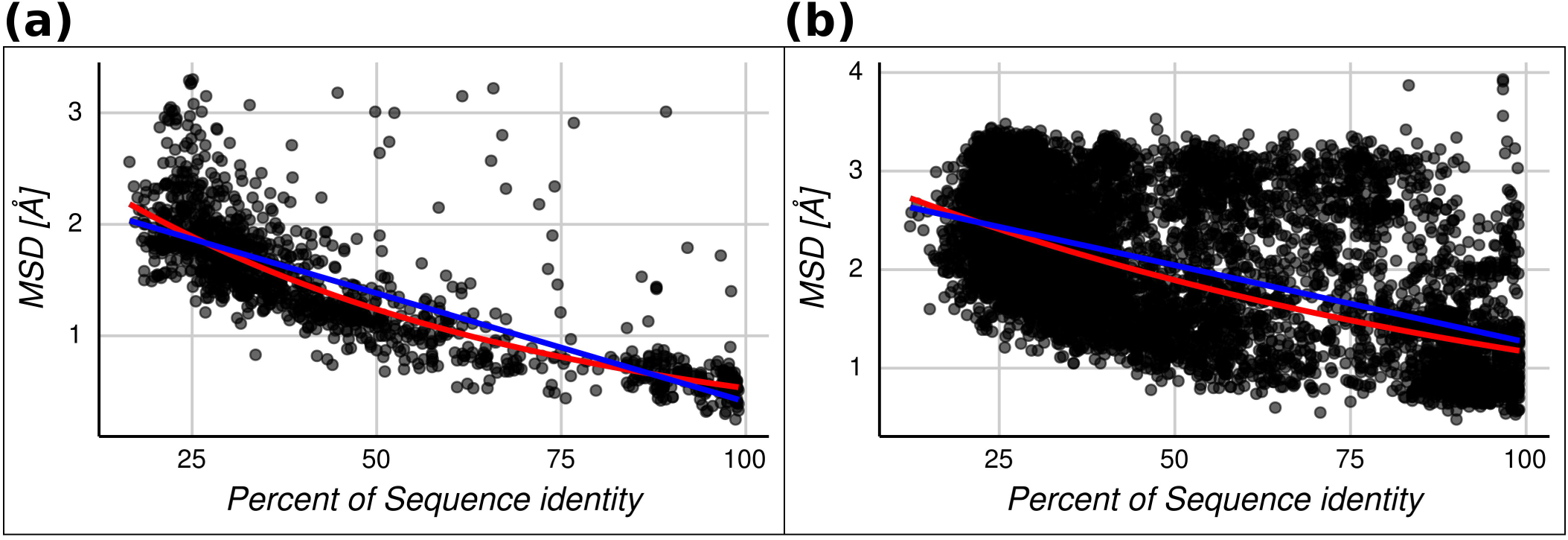
Maximum structural divergence (MSD) versus percent of sequence identity for each homologous protein pair. The lineal (blue line) and exponential (red line) regressions are shown for two sets. (a) Homologous protein pairs with an average conformational diversity less or equal than 0.5 Å. The linear and exponential fitted expressions are *RMSD* = 2.354 – 0.019 *SEQID* and *RMSD* = *e*^1.06^ *e*^−0.017 *SEQID*^, respectively. (b) Homologous protein pairs with an average conformational diversity greater than 0.5 Å. The linear and exponential fitted expressions are *RMSD* = 2.823 – 0.015 and *e*^1.121^ *e*^−0.010 *SEQID*^, respectively.

Accordingly to these results, TBM approaches will be much more reliable in protein families with low conformational diversity because the expected change in structure is proportional to the sequence divergence. In these families, where selecting the template as the one showing the highest sequence similarity and coverage, will increase the reliability of TBM. As we can see in **Fig 5a**, both linear and exponential regressions give a RMSD ~0.45 for 100% sequence identity. On the contrary, in families with a larger CD, that relationship loses predictability due to the observed structural variability for the same sequence (~1.3 for both linear and exponential regressions at 100% sequence identity).

How can we turn these findings into practical advice for use in TBM methods? It is very difficult to know the conformational diversity of the target sequence to be modelled by TBM protocols before starting. However, our previous work shows that proteins with disordered regions have larger conformational diversity compared with ordered proteins, on average[36]. In the next section, we address this question: How is protein disorder related to structural divergence?

### How does protein disorder correlate with structural divergence?

Disordered regions in proteins are known to be involved in several important biological functions[37, 38]. Intrinsically disordered regions (IDRs) or proteins (IDPs) are characterized by their high flexibility and mobility, displayed as missing regions in crystallographic structures[39]. It is difficult to estimate the extent of flexibility in the disordered regions, but it is possible to measure conformational diversity in the ordered regions of these proteins using the RMSD between different conformers, for example[40]. We found that proteins with IDRs have larger conformational diversity than those with ordered structures, when disorder-order transitions take place between protein conformations[36]. Furthermore, we recently found that proteins with IDRs can be split into two groups with different structure-function relationships, depending on how structure-based features change among the available conformer population for each protein[41]. Therefore, it is interesting to ask if the structure-sequence relationship could be also separated into two groups, mainly proteins with and without IDRs, following the above-mentioned results using CD. We find that pairs of homologous proteins containing at least one disordered region (in any of their available conformers) show higher values of MSD than the population of ordered homologous proteins (see **Fig 6**). These distributions were found to be statistically different using the Wilcoxon and Kolmogorov Smirnov test with P-value < 0.01.

**Fig 6.**
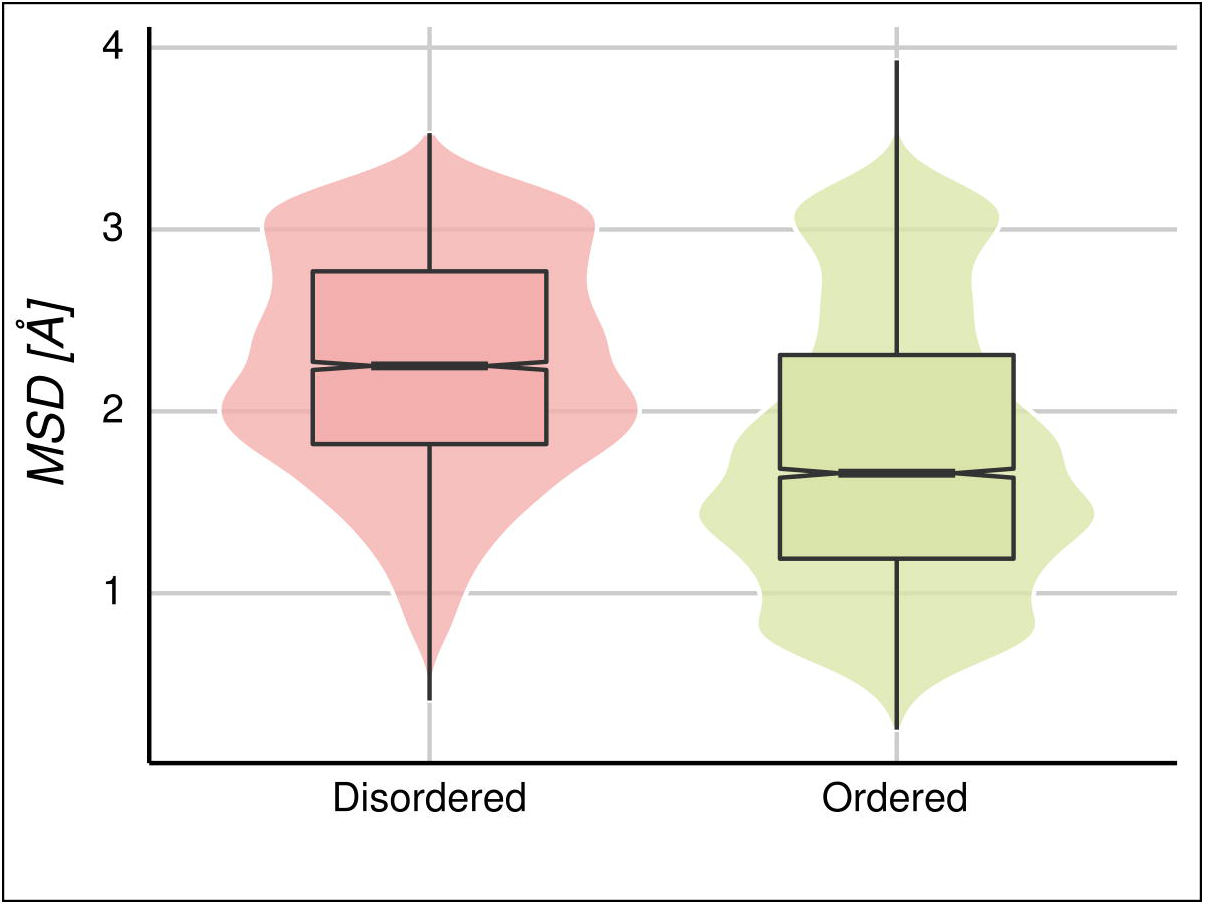
MSD distributions in homologous protein pairs. The disordered set (4439 homologous protein pairs) has proteins with at least one conformer with IDRs, while the ordered set (5034 homologous protein pairs) has no proteins with IDRs, in any of the conformers.

Since homologous pairs containing disordered regions have higher MSDs, we expect that the correlation between structure and identity percent would be worse, as shown above (**Fig 5**). We found that the Spearman’s rank correlation rho is -0.36 and -0.58 for disordered and ordered pairs of homologous proteins, respectively. These results show that the presence of disordered regions in the template and/or in the target sequence could predict a large CD and could make the relationship between sequence and structure less predictable. Although this high correlation value between full ordered proteins and MSD, using both linear and exponential fits still produces high MSD dispersions at high percent identity (**S6 Fig**). However, we know that there are few full-ordered protein families with very large CDs. These proteins have been extensively studied thanks to the pioneering work of Chothia, Lesk and Gerstein[20,21,42] and represent less than ~20% of our dataset[41]. Removing these highly dynamic proteins with non-disordered regions, we obtain the relationship shown in **Fig 7** where the Spearman’s rank correlation rho is -0.71 for the ordered set of pairs.

**Fig 7.**
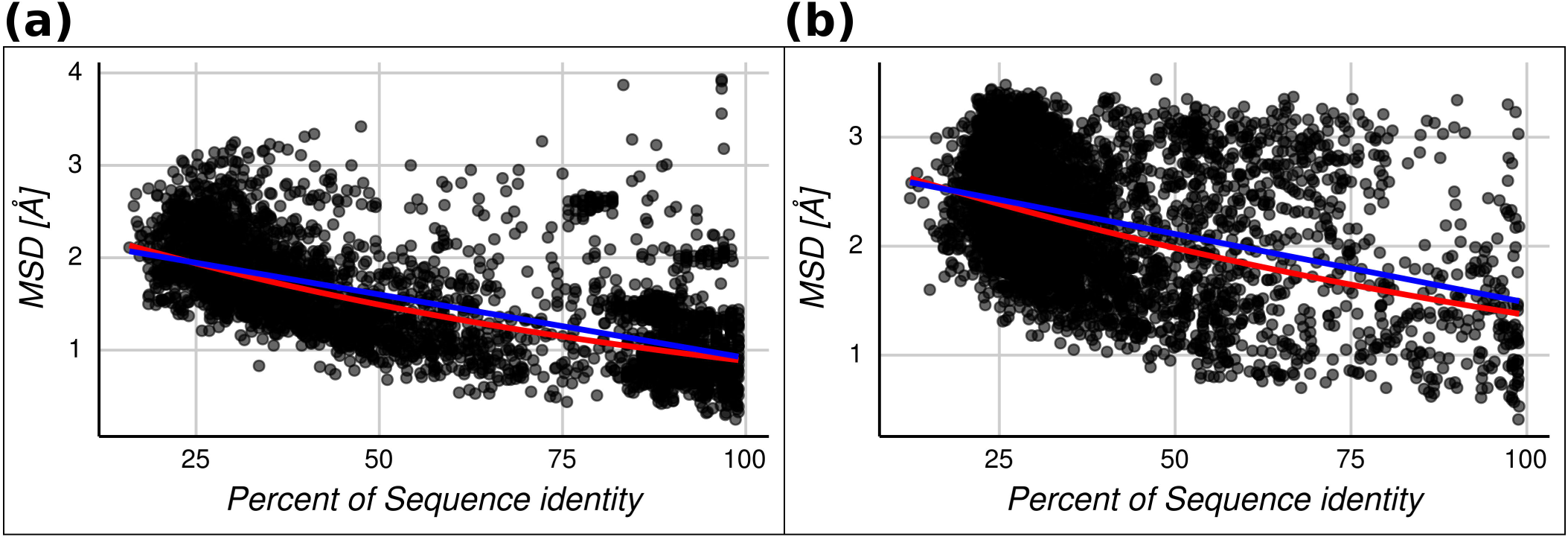
Maximum structural divergence (MSD) versus percent of sequence identity for each homologous protein pair. The lineal (blue line) and exponential (red line) regressions are shown for two sets. (a) Homologous protein pairs containing just ordered conformers. The linear and exponential fitted expressions are *RMSD* = 2.288 – 0.014 *SEQID* and *RMSD* = *e*^0.092^ *e*^−0.01 *SEQID*^, respectively. (b) Homologous protein pairs with at least one of the conformers containing disordered regions. The linear and exponential fitted expressions are *RMSD* = 2.741 – 0.012 *SEQID* and *RMSD* = *e*^1.055^ *e*^−0.007 *SEQID*^, respectively.

Based on these correlation coefficients, we can say that the presence of disorder regions alone has a moderate capacity for predicting the presence of noise between structure and sequence variables. Moreover, these values were obtained considering that most ordered proteins do not have CD, or at least show moderate CD, as we have previously described[41]. Taking into account these considerations, and the easy and reliable prediction capacity for disordered regions in proteins, this information could be still used as guidance in TBM approaches.

## Discussion

The study of structure-sequence relationships embraces a foundational concept for several areas focused on the study of proteins. Biological function prediction[43], protein evolution[44], structural proteomics[45], and homology modeling[3] are just a few examples of the broad and active research areas that take advantage of that relationship. Among all the situations that structure-sequence could adopt, we have focused on how the presence of conformational diversity in proteins could influence the relationship between sequence and structural change and therefore affect TBM approaches. A central point was derived early by Chothia and Lesk, showing that the success of 3D prediction will depend of the extent of the target sequence identity with the corresponding template[7]. Basically, the behaviour between RMSD and percent identity established a relationship in which structural similarity increased as sequence similarity increased. Their results, and conclusions, have been verified by numerous studies[8,9,11,12,46–48]. These studies found moderate-to-high correlation coefficients between different parameters that were proportional to structural and sequence similarity, i.e. RMSD and percent identity[9, 48], evolutionary distance[12], and statistical significance of RMSD[13]. They also found linear and nonlinear behaviour, and an invariably low structural variation, at 100% identity (~0.5 Å). Departure from linear fitness has been explained as being derived from errors in the alignments (at high and low sequence similarity), use of redundant datasets, or by ignoring the multiple substitutions per site during evolution[12,13,46].

In this work, we found that the extent of the CD is related to the MSD of the family, and that the structure-sequence relationship is more complex than previously thought. First, we found that the extent of the CD could be as large as the MSD (**Fig 1**). Conformational diversity is a key concept to understand many processes and mechanisms in protein function, such as enzyme catalysis[49], promiscuity in protein interactions[50], protein-protein recognition[51], signal transduction[52], mechanisms of disease-related mutations[53], immune escape[54], the origin of neurodegenerative diseases[55], protein evolutionary rates[56], conformer-specific substitution patterns[57], the origins of new biological functions[58], molecular motors[59], and co-evolutionary measurements between residues[24, 60] (for a recent review please see[17]). Furthermore, we have recently shown that the distribution of CD in a large dataset of proteins, with experimentally determined CD (~5000), results in three main groups of proteins with different structure-function relationships[41]. Distributions in **Fig 1** show that CD could be as large as the MSD, but it is also evident that most of the proteins in our dataset have modest to low CD, meaning that they could function with very low or absent backbone movements[41,61,62].

Several studies drew attention to the importance of CD in TBM methods[18,19,26], but the consideration of CD in the study of structure-sequence relationships, as an essential ingredient in TBM methods, was often considered a source of bias or “noise”[12, 13]. On one side, avoiding the “noise” introduced by the use of redundant data (i.e. considering CD) would allow us to assume that structural changes would be proportional to sequences changes. The existence of a single sequence with multiple conformations defines a “degeneracy” of the structural information coded in a given sequence, introducing a nonlinear behavior in the protein space. It was also found that this nonlinearity could possibly impair the performance of knowledge-based methods in bioinformatics[63].

However, taking into account the remarkable importance of CD for explaining biological processes and protein behavior, it appears impossible to ignore. As derived from **Fig 2**, when expressed as an average between all families, there is a large dispersion in RMSD even at high sequence identities but interestingly, this dispersion remains at the family level (**Fig 3**). These figures show the large uncertainty in template selection even at high sequence similarities. We have also found that CD is proportional to the MSD reached in a family (**Fig 4**). Indeed, CD is independent of the maximum sequence divergence of the family (**S5 Fig**). It is at this point that the total set used in this work could be split in two sets of reduced and large CD impact. In proteins evolving under selective pressure to maintain a reduced CD, we find that the correlation between sequence and structure variation is high (Spearman’s rank correlation rho = -0.83). We have previously characterized this group of proteins[41]. Briefly, rigid proteins have low CD (~0.8 Å RMSD in average), they are mainly proteins without disordered regions and have important tunnels and cavities. When conformers of rigid proteins are compared, for example in their bound and unbound states, we find that they mainly differ in backbone positions associated with their tunnels and cavities. This indicates that the minimum movements for rigid proteins are associated with the movement of functional structures to allow the transit of substrates and/or products between the inside and the surface of the protein[64–66]. It is for this group of proteins that sequence-structure relationships show a high correlation between variables, and for whom it would be possible to reliably predict 3D models using TMB techniques (see **Fig 5a**).

According to our results, proteins with higher CD also have larger structural divergences (**Fig 4**), and a larger variability of accessible structures, even at high percentages of sequence identity (**Fig 5b**). We found that correlations and proportionalities between variables in this group of proteins are low, blurring the basic idea common in TBM approaches that similar sequences show similar structures. For example the expected value at 100% identity using the linear regression gives ~1.3 Å.

Our results indicate that the relative success of the 3D model using TBM approaches will be strongly associated with the CD of the corresponding target protein. Because it is difficult to predict the extent of CD in a given protein, we used the presence of disordered regions as an indicator of higher values of CD, based upon previous results[36]. We found that the presence of highly ordered proteins (without any disordered regions in any conformer) in pairs of homologous proteins have a Spearman’s correlation rho of -0.58 for RMSD and a sequence identity relationship. In this way we found that presence of disorder/order is not a strong indicator of a well-correlated sequence and structural change (**Fig 7**), compared with the knowledge of the extent of CD (Spearman’s correlation rho of -0.83, **Fig 5**). Removing ordered and highly dynamic proteins, we found a better Spearman’s correlation rho of -0.71. This increment in reliability again confirms the higher correlation between sequence and SD for rigid proteins with low CD. Based on the many and reliable predictor methods for detecting disordered regions in proteins[67], and the above mentioned considerations, order/disorder could be an easy way for evaluating the expected dispersion of RMSD for a given sequence similarity between template and target sequences. Alternatively, since the CD of some proteins is well correlated with the MSD of the family (**Fig 4**), comparing all the known structures of the family can predict the expected flexibility of our target. However, further studies and experimental data are required in order to address the question of how well CD is conserved through evolution.

In summary, sequence and structure divergence is a more complex process than previously thought. Protein conformational diversity challenges the ordered and well-accepted relationship between sequence and structural similarity, a cornerstone of TBM techniques, as well as our understanding of the nature of the protein folding code. Further work is necessary to deepen our current knowledge in such a basic topic for many areas associated with the study of proteins, as well as to encourage a reappraisal of current methods for obtaining and evaluating 3D protein models.

## Methods

### Protein families with conformational diversity selection

The CoDNaS database[23], containing a redundant collection of three dimensional structures for the same protein (at least 95% of sequence identity among structures to include putative sequence variations), was used to recruit proteins exhibiting conformational diversity. All structures belonging to this dataset were obtained by X-Ray Diffraction at a resolution equal or less than 2.5 Å. The maximum conformational diversity for each protein (CD) is the maximum C-alpha RMSD derived from all conformer pairwise comparisons. With the aim to obtain a reliable and comparable estimation of conformational diversity of each protein, our dataset only contains proteins with a minimum of 5 conformers (average ~19 conformers per protein) as was previously suggested[68].

In order to identify homologous proteins, we ran BLASTClust[69] to obtain all available clusters at 30% of local sequence identity with a minimum coverage of 0.9 between all sequences in the cluster. The PDB SEQRES records were used to extract the sequences and to perform the clustering. We only considered those clusters with at least two different proteins using the UniProt ID for identifying each protein.

The final dataset contains 2024 different protein chains with a total of 37755 conformers. These proteins are grouped in 524 families with an average of ~4 proteins per family (with a minimum of 2 and a maximum of 61).

### Sequence and structure comparisons

To estimate the SD for each homologous protein pairs in a cluster, we calculated the C-alpha RMSD using MAMMOTH[27] for all possible pairs of conformers belonging to the proteins being compared. MAMMOTH is a sequence-independent structural alignment program, which not only has a very good accuracy aligning proteins with different folds, but also provided the statistical reliability of the resulted structural alignment. Also, the RMSD values calculated by MAMMOTH show no dependence with protein size nor length.

The MSD for a pair of homologous proteins is the pair with maximum RMSD value among all vs all conformers comparisons between them. Additionally, we calculated the percent sequence identity for each homologous protein pairs using a global sequence alignment obtained with the Needleman-Wunch algorithm[28]. Furthermore, we defined disordered regions for each conformer when it has five or more consecutive missing residues that were not in the amino or carboxyl terminal of the protein sequence (the first or least twenty residues). If a residue has missing electron density coordinates in a structure obtained with X-ray crystallography, it assumed to be disordered[70]. If a protein has at least one conformer with disordered regions we classified it as disordered.

The total comparisons among all vs all conformers for each homologous protein pairs and structures of the same protein give an amount of ~3.5 million of pairs.

### Statistical analysis

The correlations coefficients showed in this work were obtained using the function *cor.test* of R package[71], with the corresponding two-way test (null hypothesis is that the correlation coefficient is equal to 0). We used Spearman rank correlations coefficients since it does not assume linearity of the data, only searches for a monotonic relationship.

Cross-validated function fitting were performed using the Julia language libraries LsqFit and MLBase. All the regressions were weighted, and each point (MSD vs sequence identity percent) has a weight of 1 over the number of protein pairs in the protein family. It was done in order to avoid that populated families predominate the results. However, the R^2^ informed was calculated as the mean of the R^2^ in each testing subset from a 5-fold cross-validation without using weights. In that way, we use the amount of unweighted variation explained by the model as a measure of goodness of fit.

## Author Contributions

Conceived and designed the experiments: AMM, CMB and GP. Performed the experiments: AMM. Analysed the data: AMM, GP, CMB and DZ. Contributed reagents/materials/analysis tools: AMM and DZ. Wrote the paper: GP, CMB, AMM and DZ.

## Supporting information

**S1 Table. Spearman’s rho coefficient and coefficient of determination (R^2^) calculated in bins of CD.**

**S1 Fig. Schematic representation of the derivation of MSD (Maximum Structural Divergence) and CD (conformational diversity)**

**S2 Fig. RMSD vs percent of sequence identity**. RMSD obtained from an all vs all comparison between two homologous proteins considering all their conformers. The figure contains about 3.5 million comparisons.

**S3 Fig. Distributions of MSD in bins of homologous protein pairs number contained in each family.** It is possible to see that the distribution of MSD is not influenced by how populated the family is.

**S4 Fig. Distribution of the standard deviation of RMSD due to CD per family estimated over the CD for each protein by family.** Standard deviations have been accumulated in two extreme cases, families with low (less than 0.5 Å of RMSD) and high CD (more than 0.5 Å of RMSD).

**S5 Fig. Relationship between MSD and CD in bins of sequence identity percent.** Each dot represents the average RMSD values for the MSD and the CD in a specific family. The number of families in each bin are 348, 138 and 38 with Pearson’s correlation coefficients of 0.77, 0.87 and 0.88 respectively.

**S6 Fig. Maximum structural divergence (MSD) versus percent of sequence identity for each homologous protein pair**. The lineal (blue line) and exponential (red line) regressions are shown. All homologous protein pairs containing just ordered conformers. The linear and exponential fitted expressions are RMSD = 2.57 - 0.015 SEQID and RMSD=e1.03 e(-0.01 SEQID) respectively.

